# DIPLOMAT: multi-animal tracking with efficient manual editing

**DOI:** 10.1101/2025.08.11.669786

**Authors:** Isaac Robinson, George Glidden-Handgis, Neekesh Panchal, Nathan Insel, Travis J. Wheeler

## Abstract

Recent advances in computer vision have enabled the development of automated animal behavior observation tools. Several software packages currently exist for concurrently tracking pose in multiple animals; however, existing tools still face challenges in maintaining animal identities across frames and can demand extensive human oversight and editing. Here we report on DIPLOMAT, a <u>D</u>eep learning-based, <u>I</u>dentity-<u>P</u>reserving, <u>L</u>abeled-<u>O</u>bject <u>M</u>ulti-<u>A</u>nimal <u>T</u>racker, which implements automated algorithms improving tolerance to occlusion and continuity of animal identity over a video, further supplemented by an efficient human interface to help eliminate remaining errors. DIPLOMAT is designed to perform multi-animal tracking by building on the per-frame pose prediction models of two state-of-the-art tools, DeepLabCut and SLEAP. Where other tools immediately take a maximum likelihood estimate from a given video frame, DIPLOMAT splits the probability fields according to the number of tracked animals and then applies an inference method that takes into account across-frame movement and probabilistic distances between body parts. These independent trace probabilities are then preserved for human editing, enabling multiple body parts to be re-tracked across frames with minimal user action. Testing with a standardized and independently tracked dataset of 3-mouse videos shows DIPLOMAT’s automated components alone can reduce identity swaps by *>*75%. DIPLOMAT code and documentation are available at https://diplomattrack.org/.

**Significance Statement:** Tracking multiple animals during social behavior is essential for understanding neural mechanisms underlying social interaction, yet current tools frequently lose individual identities when animals approach/overlap. This identity confusion can require users to spend hours manually correcting tracking errors. DIPLOMAT addresses this bottleneck by integrating neural network and HMM approaches, combining per-frame body-part probabilities with probabilities related to inter-body-part distances and movement through space. We find this reduces identity swap errors without sacrificing other metrics. Moreover, DIPLOMAT stores probability data for use in a manual editing tool that allows for multi-body-part, manual re-tracking and error correction. DIPLOMAT therefore reduces user-time requirements for processing multi-animal recordings, which has the potential to accelerate research into the neural and genetic basis of social behavior.

## Introduction

During social behavior, one individual’s movements can depend on subtle, moment-to-moment changes in another’s facial expressions or posture. Our ability to fully understand social interaction – from biological mechanism to its role in health or ecology – may depend on methods to monitor the complexity of these movements across multiple animals. Computer vision advances have made this increasingly possible, but there are still many practical limitations. A particular challenge has been in keeping track of identities: where standard algorithms excel at identifying the locations of body parts in a given image, they often still falter in maintaining a consistent identity for the owner of those parts across a recorded video. The fundamental issue is that the ability to universally (abstractly) recognize a body part of any individual can be contradictory to the ability to distinguish between individuals.

Several tools for multi-animal tracking have recently become available, each with slightly different approaches toward maintaining individual identities. This includes ID tracker [1], AlphaTracker [2], TRex [3], SLEAP [4], multi-animal DeepLabCut (DLC; [5]), and PrecisionTrack [6]. Three of these, DLC, SLEAP, and AlphaTracker have caught-on as particularly popular and powerful tools for tracking dyads (two individuals) and triads (three individuals) of rodents using a single-camera video source [7–10]. Each of these three tools assigns identities on individual frames, then stitches them together across frames using various similarity-matching methods. For assignments within each frame, DLC follows a “bottom-up” (parts-to-bodies) approach, predicting probability distributions for body parts then assembling them into per-animal skeletons (whole bodies) via graph/geometry constraints. AlphaTracker follows a “top-down” (bodies-to-parts) approach, first detecting animal bounding boxes, then identifying body parts within each box as belonging to a single animal. SLEAP supports both bottom-up and top-down part assignments. To stitch identities across frames, the tools use different but related methods: DLC uses spatial proximity and pose similarity, AlphaTracker uses overlap of bounding box (body) and probability distribution (body parts), and SLEAP likewise uses pose-space similarity along with motion models (optical-flow/Kalman style predictions) to account for frame-to-frame displacement. PrecisionTrack uses a Dynamic Kalman Filter as the motion model. In cases where tracked bodies are visually distinct, additional algorithms can be applied; for example, in DLC, users can specify identifying features, or use a transformer-based “reID” method that automatically detects visual distinctions.

Each of the above approaches to maintaining animal identity from one frame to the next is theoretically viable; empirically, however, they are still known to produce cases in which individual body parts or entire skeletons are swapped between individuals. Errors are particularly common when animals are near or on top of one-another, causing full body swaps or skeletons spanning both bodies (“chimeras”). While post-tracking user-edits can help correct for these, editing can be laborious. In SLEAP, for example, correcting a skeleton that spans both bodies requires clicking and dragging each individual point from one body to the other–for each frame that they are mislabeled (even if the error lasts only one second of video time, at 60 frames/s that can require moving multiple body parts for each of 60 frames).

There are therefore two challenges with current, 2-D tracking methods: First, they tend to make many identity errors – of single body parts or of entire bodies – when there are similar-looking animals that sometimes occupy overlapping or nearby spaces. Second, tools for manually correcting tracking errors are laborious and inefficient.

Here, we introduce DIPLOMAT (<u>D</u>eep learning-based, <u>I</u>dentity-<u>P</u>reserving, <u>L</u>abeled-<u>O</u>bject <u>M</u>ulti-<u>A</u>nimal <u>T</u>racker), which seeks to improve identity accuracy and persistence in multi-animal tracking. Rather than introducing an entirely new tool, we designed DIPLOMAT as a method capable of leveraging and expanding on the effective per-frame pose prediction results of other tools. DIPLOMAT currently uses models trained by either DLC or SLEAP as the basis of per-frame pose prediction, then applies a distinct set of downstream algorithms to make predictions about body part location and animal identity. Importantly: the goal of DIPLOMAT is not to address all possible multi-object tracking options, but to specifically introduce algorithms and editing tools that improve identity preservation independent of visual distinctions between objects (i.e., can be applied to tracking nearly identical animals). We also treat characterization of social interaction based on multi-animal pose estimation as outside of the scope of the software; other tools that address automated social classification, primarily in mice, include JAABA [11], MARS [12], MoSeq [13], and SimBA [14].

DIPLOMAT is developed with two components in mind. The first (`Track`) implements algorithms that help automatically track body parts of multiple animals while minimizing identity errors. The second (`Interact`) employs an intuitive and efficient user interface for rapid editing of multiple body parts across video frames, with the ability to re-integrate these edits to a quickly and smoothly re-track.

DIPLOMAT casts the tracking problem as one in which bodies and their parts have been produced by a Markov process – in which state at a given time depends on the state at nearby times – and can thus be addressed with a hidden Markov model (HMM). DIPLOMAT treats the per-frame pose labeling confidence values from neural-network models (trained using DLC or SLEAP) as the source of the Markov model observation probabilities. It then applies the HMM Viterbi algorithm [15] to identify a maximum probability trace across the full history of frames in a video. To compute transition probabilities for the Markov model, both body motion and body part (skeletal) connections are taken into account. To take into account multiple, similar bodies (multiple animals), an additional step is taken that splits probability fields in each frame, generating what we call “mutually independent trace”. The full-video perspective of the Viterbi algorithm allows DIPLOMAT to avoid making identity switch errors that may occur using methods that only focus on stitching identity between neighboring frames.

DIPLOMAT methods are guided by 4 assumptions: **(1) Bodies don’t teleport:** neighboring video frames (past and future) can be used to inform probability distributions for current body part locations. **(2) Body parts stick together:** skeletal information can further inform a posterior probability of body part location. **(3) Bodies will be easier to distinguish in some frames:** some frames can serve as an anchor, or “ground truth”, and those anchored assignments can influence the labeling of neighboring frames. Finally, **(4) in a given recording, there is typically a pre-set number of bodies:** probability distributions can be sliced into a fixed number of independent probability clusters. Although not all of these assumptions will be true for every application, they cover a wide scope of animal behavior and neuroscience protocols.

In this paper we provide a detailed description of the algorithms supporting DIPLOMAT tracking, and demonstrate that DIPLOMAT’s automated methods reduce identity-swaps in a standard benchmarks dataset (MABe; [16]). We further describe a novel *Interact* interface that allows users to dynamically and efficiently correct inferred body part locations in frames. A key feature of Interact over other manual editing tools is that it makes use of probability maps from pose estimation neural networks to facilitate fast and accurate single- or multi-body part editing. Whereas other tools discard per-frame part placement probability distributions during the course of tracking, DIPLOMAT explicitly retains them for use in semi-automated support of manual correction. Interact includes the ability to dynamically re-trace (re-run the independent-trace Viterbi algorithm) across frames, thereby incorporating user-made edits.

## Methods

DIPLOMAT addresses the challenge of identity swaps during multi-animal tracking by integrating history and relational information relevant for identity inference into the tracking process. The process consists of multiple stages. Briefly, they are:

- The user trains a neural network model capable of estimating the pixel-by-pixel probabilities that a body part is present in a video frame. Several software environments for this (pose inference engines) exist, and do not require any connection to DIPLOMAT. Currently, DIPLOMAT supports DLC and SLEAP, in part due to their popularity, and also because their bottom-up approaches allow DIPLOMAT to track using the same models.
- DIPLOMAT applies a model to a video to compute pseudo-probabilities per body part across all pixels of each video frame.
- DIPLOMAT performs a crude analysis on the per-frame pseudo-probabilities to estimate body-part distances and parse the video into segments. It clusters neighboring pixels, computes estimates of body/skeleton size(s), identifies frames where the tracked animals are particularly far apart (“anchor frames”), and splits video into time segments separated by those frames.
- For each segment, for each body part, DIPLOMAT applies the HMM Viterbi method to compute a maximum probability trace; i.e., a path through the video frames that takes into account each frame’s neural network probabilities along with transition probabilities based on estimated movement and inter-body distance. One key modification in computing the maximum probability trace is that probabilities are split according to the number of animals, so that traces for each are computed in a way that is mutually independent. Details on these calculations are provided below.
- The result is a set of mutually independent traces within each segment, one for each body part of each tracked animal. Since the within-segment computations are independent from one-another, they can be computed in parallel, optimizing runtime. DIPLOMAT then stitches the segments together, using the Hungarian algorithm, to preserve identity across segments.
- The probabilities are saved, and the methods can be re-run on-the-fly, in a user-friendly graphical interface, `Interact`.
- An additional, optional tool, `Tweak`, provides another point-and-click interface to edit single body part positions or multi-body-part ID swaps for final CSV table outputs.

A detailed description of the algorithms supporting DIPLOMAT tracking is available in the Extended Methods section near the end of the manuscript.

### Code accessibility

DIPLOMAT is open-source software released under the BSD 3-clause license. Code and documentation are available at https://diplomattrack.org/.

## Results

### Tracking process

DIPLOMAT can be installed using the DIPLOMAT PyPi package (https://pypi.org/project/diplomat-track/). To use DIPLOMAT for tracking, a user must first train a pose inference model. Currently, DIPLOMAT supports models trained using the open-source packages, DLC and SLEAP (bottom-up). Installation and training method for these tools have been detailed elsewhere [4, 5].

Given a video and a trained model, DIPLOMAT’s `Track` module performs per-frame pose inference using a custom implementation of the inference engine appropriate for the trained model. Following per-frame pose inference, DIPLOMAT seeks a trace for each body part (keypoint) that maximizes trace probability under a model that mixes pose inference probability fields with movement distance probabilities supported by skeletal influence. Following automated tracing, the user may *Interact* with the resulting trace to correct errors. See Figure 5 for an overview, and Methods for details.

### Validation

Quality metrics were computed for DIPLOMAT in conjunction with the standalone SLEAP and DLC. As a benchmark, we used the MABe “Mouse triplet” data set as a source of ground-truth standardized videos [16]. The MABe dataset is one-of-a-kind for multi-animal validation because of how the ground-truth locations were determined: each body part of each frame of each video was annotated independently by 5 human workers. Thus, locations were identified entirely independently from DeepLabCut, SLEAP, or DIPLOMAT, ensuring that no tool had a relative advantage. Fifteen, 60-second videos from MABe were used to train models (27,000 frames). Training and testing were performed using a system with two Xeon E5-2695v4 14-core processors, 192GB system RAM, and two Nvidia P100 GPUs with 16GB RAM.

Tracking was performed using each tracking tool, across the 60-second evaluation MABe mouse triplet videos, filtering to retain only the 54 videos that contained position data for each body part of interest for each frame. Tools were tested under two body architectures: one included 10 body parts (nose, ears, neck, forepaws, hind paws, center-back, and tail base) and the other only 5 body parts (nose, ears, center-back, tail-base). We refer to these as BPM-10 and BPM-5 respectively. Metrics were computed using the MotMetrics analysis package [17]; in particular, body swaps are computed using the *num switches* metric. The set of videos proved to be an adequate sample to detect differences between software tools, as over 40% of videos (24 and 23 for BPM-10 and BPM-5 respectively) induced at least one tracker to produce at least one “full body swap” (i.e., where most body parts of an animal moved to another body), while over 70% (39 and 40 in for BPM-10 and BPM-5 respectively) induced at least one body part swap in at least one tracker.

DIPLOMAT improved tracking relative to both DLC and SLEAP when using either BPM-10 or BPM-5. Using the BPM-10, the incidence of body-swaps across individual body parts was lower when DIPLOMAT was used against DLC or SLEAP (Figure 2 A&B; top row of Tables 1; Wilcoxon signed rank test for zero median). Strong reductions were also observed in full body swaps (Figure 2C&D; top row of Table 2). Using BPM-5, body part identity swaps were reduced compared to stand-alone DLC (Table 1), and full body swaps reduced relative to SLEAP, with a strong statistical trend against DLC (Table 1). DIPLOMAT also improved scores across other important measures. ID recall – the proportion of body parts (Table 1) and bodies (Table 2) that DIPLOMAT correctly identified – was consistently higher with DIPLOMAT. DIPLOMAT also improved whole-body ID precision, a measure inversely related to the number of false-positives, and for individual body parts showed improvements over SLEAP but not DLC. Finally, DIPLOMAT tended to show better proximity of points identified relative to ground-truth points (MOTP) when compared against the same model used in stand-alone SLEAP, with only minor differences observed compared to the same stand-alone DLC model.

**Table 1.**
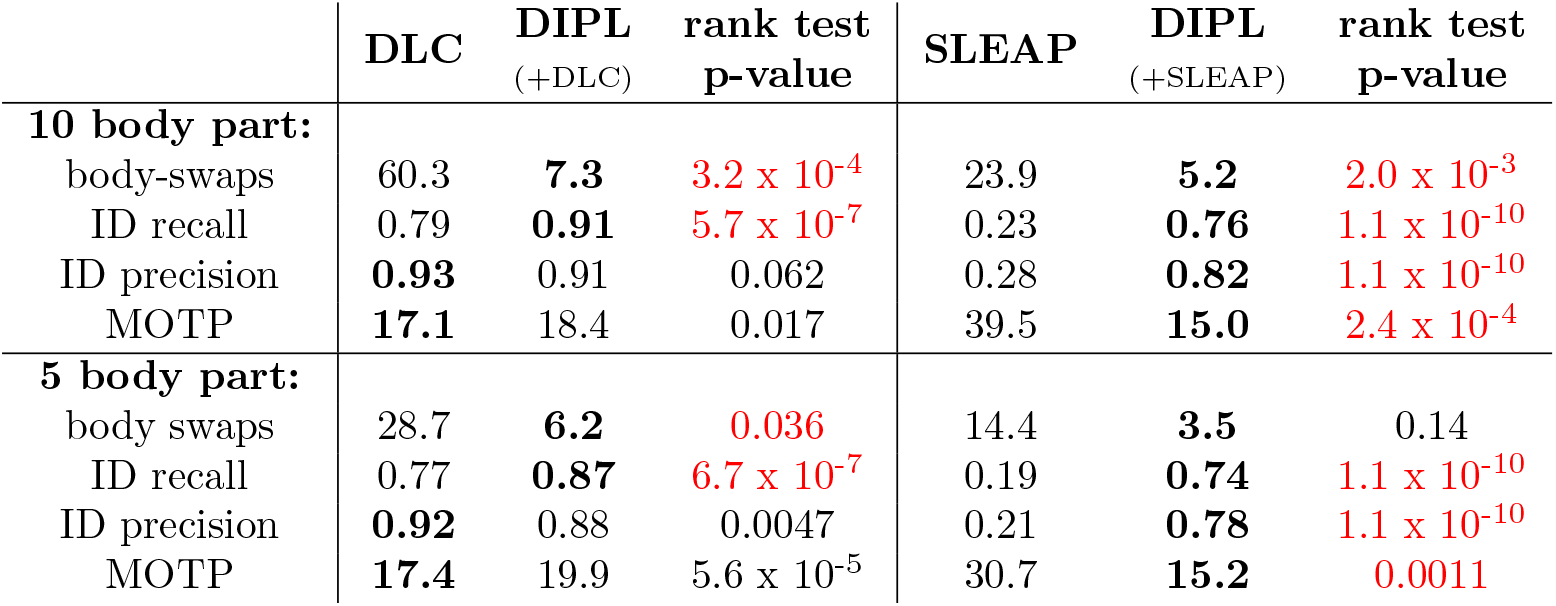
Means and Wilcoxon signed-rank test p-values across all individual body parts. DIPL(+DLC) represents results using DIPLOMAT on top of DLC per-frame inferences, while DIPL(+SLEAP) uses SLEAP inferences. Bold text indicates the superior outcome in each pairwise comparison. Red text indicates significant (P *≤*0.05) improvement with DIPLOMAT.

|  | DLC | DIPL<br>(+DLC) | rank test<br>p-value | SLEAP | DIPL<br>(+SLEAP) | rank test<br>p-value |
| --- | --- | --- | --- | --- | --- | --- |
| **10 body part:** |  |  |  |  |  |  |
| body-swaps | 60.3 | **7.3** | $3.2 \times 10^{-4}$ | 23.9 | **5.2** | $2.0 \times 10^{-3}$ |
| ID recall | 0.79 | **0.91** | $5.7 \times 10^{-7}$ | 0.23 | **0.76** | $1.1 \times 10^{-10}$ |
| ID precision | **0.93** | 0.91 | 0.062 | 0.28 | **0.82** | $1.1 \times 10^{-10}$ |
| MOTP | **17.1** | 18.4 | 0.017 | 39.5 | **15.0** | $2.4 \times 10^{-4}$ |
| **5 body part:** |  |  |  |  |  |  |
| body swaps | 28.7 | **6.2** | $0.036$ | 14.4 | **3.5** | 0.14 |
| ID recall | 0.77 | **0.87** | $6.7 \times 10^{-7}$ | 0.19 | **0.74** | $1.1 \times 10^{-10}$ |
| ID precision | **0.92** | 0.88 | 0.0047 | 0.21 | **0.78** | $1.1 \times 10^{-10}$ |
| MOTP | **17.4** | 19.9 | $5.6 \times 10^{-5}$ | 30.7 | **15.2** | $0.0011$ |

**Table 2.** Means and Wilcoxon signed-rank test p-values for whole body (i.e., mean of all body parts). Headers as in the previous table. Bold text indicates the superior outcome in each pairwise comparison. Red text indicates significant (P *≤*0.05) improvement with DIPLOMAT.

|  | DLC | DIPL<br>(+DLC) | rank test<br>p-value | SLEAP | DIPL<br>(+SLEAP) | rank test<br>p-value |
| --- | --- | --- | --- | --- | --- | --- |
| **10 body part:** |  |  |  |  |  |  |
| body swaps | 1.8 | **0.29** | $0.044$ | 3.8 | **0.4** | $0.0035$ |
| ID recall | 0.86 | **0.98** | $5.8 \times 10^{-6}$ | 0.89 | **0.93** | $0.0085$ |
| ID precision | 0.87 | **0.98** | $5.8 \times 10^{-6}$ | 0.89 | **0.93** | $0.018$ |
| MOTP | **15.4** | 17.2 | 0.46 | 58.6 | **40.5** | $1.8 \times 10^{-4}$ |
| **5 body part:** |  |  |  |  |  |  |
| body swaps | 1.6 | **0.38** | 0.052 | 4.7 | **0.56** | $0.0015$ |
| ID recall | .84 | **.96** | $4.5 \times 10^{-7}$ | 0.71 | **0.83** | $5.5 \times 10^{-4}$ |
| ID precision | 0.84 | **0.96** | $4.5 \times 10^{-7}$ | 0.73 | **0.82** | $0.0045$ |
| MOTP | **14.7** | 21.0 | 0.24 | 170 | **38.6** | $6.0 \times 10^{-10}$ |

**Fig 1.**
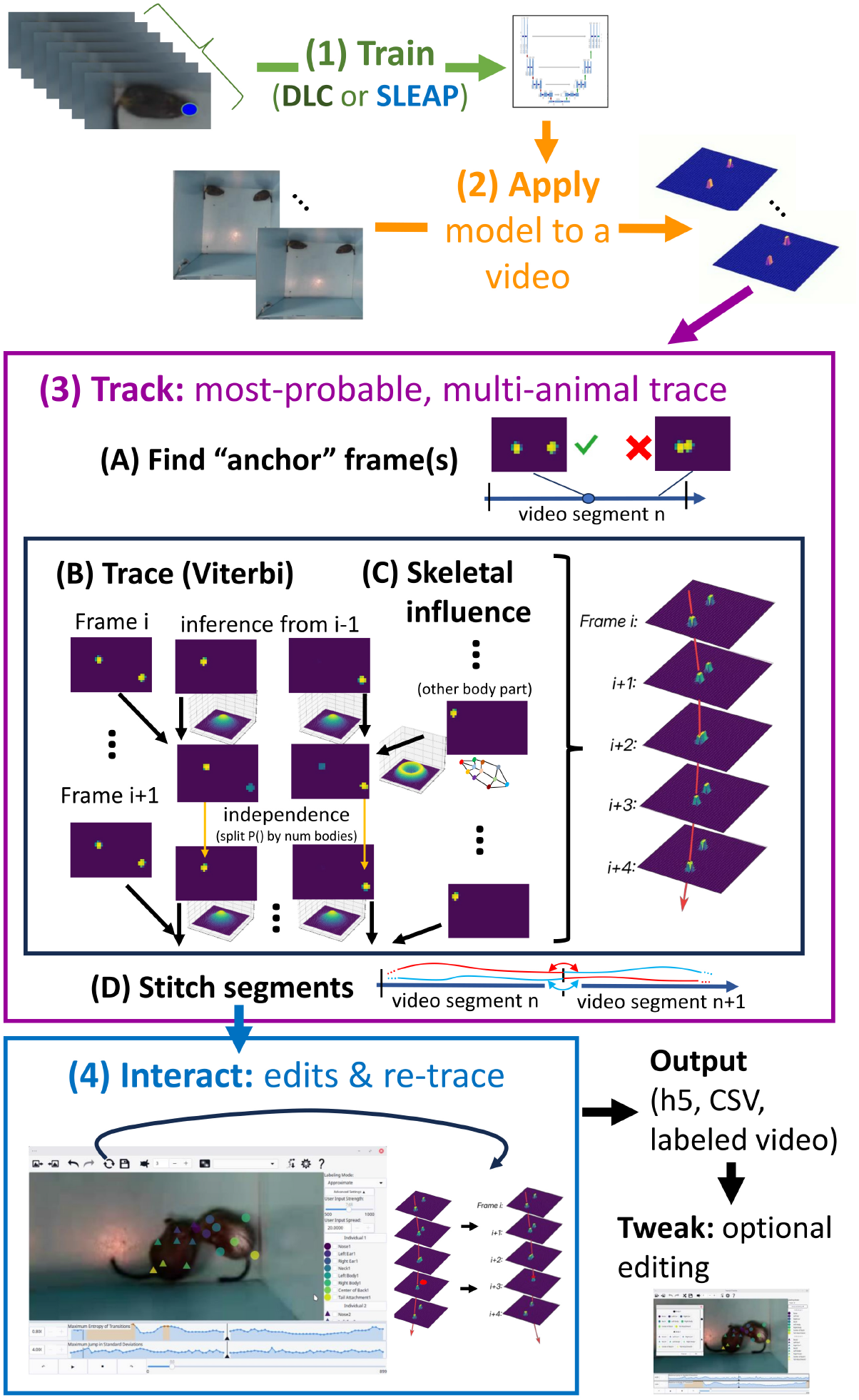
Flowchart illustrating the sequence of steps involved in multi-animal tracking using DIPLOMAT. Step 1 (top): Either SLEAP or DLC is used to train a model by manually labeling frames. Step 2: DIPLOMAT applies SLEAP or DLC inference functions to generate frame-by-frame probabilities on a video. Step 3: DIPLOMAT segments the video by identifying “anchor-frame” locations where animals are spatially separated (A), then combines a Viterbi trace (B) with skeletal constraints (C) to generate a map of transition probabilities. For each frame, independence between tracks for different individuals is imposed by splitting transition probabilities based on the most prominent path (dominance). Segments are then stitched together (D), using a Hungarian algorithm, which maps probability peaks, to preserve identity between segments. Step 4: Errors in tracking can be identified and corrected using the Interact interface, which leverages the same probability maps to assist with single- or multi-body part edits, and allows the automated algorithms to be re-run. The output of Interact is a table of x-y position data across all body parts and individuals. These can be further edited using a paired-down interface `Tweak.`

**Fig 2.**
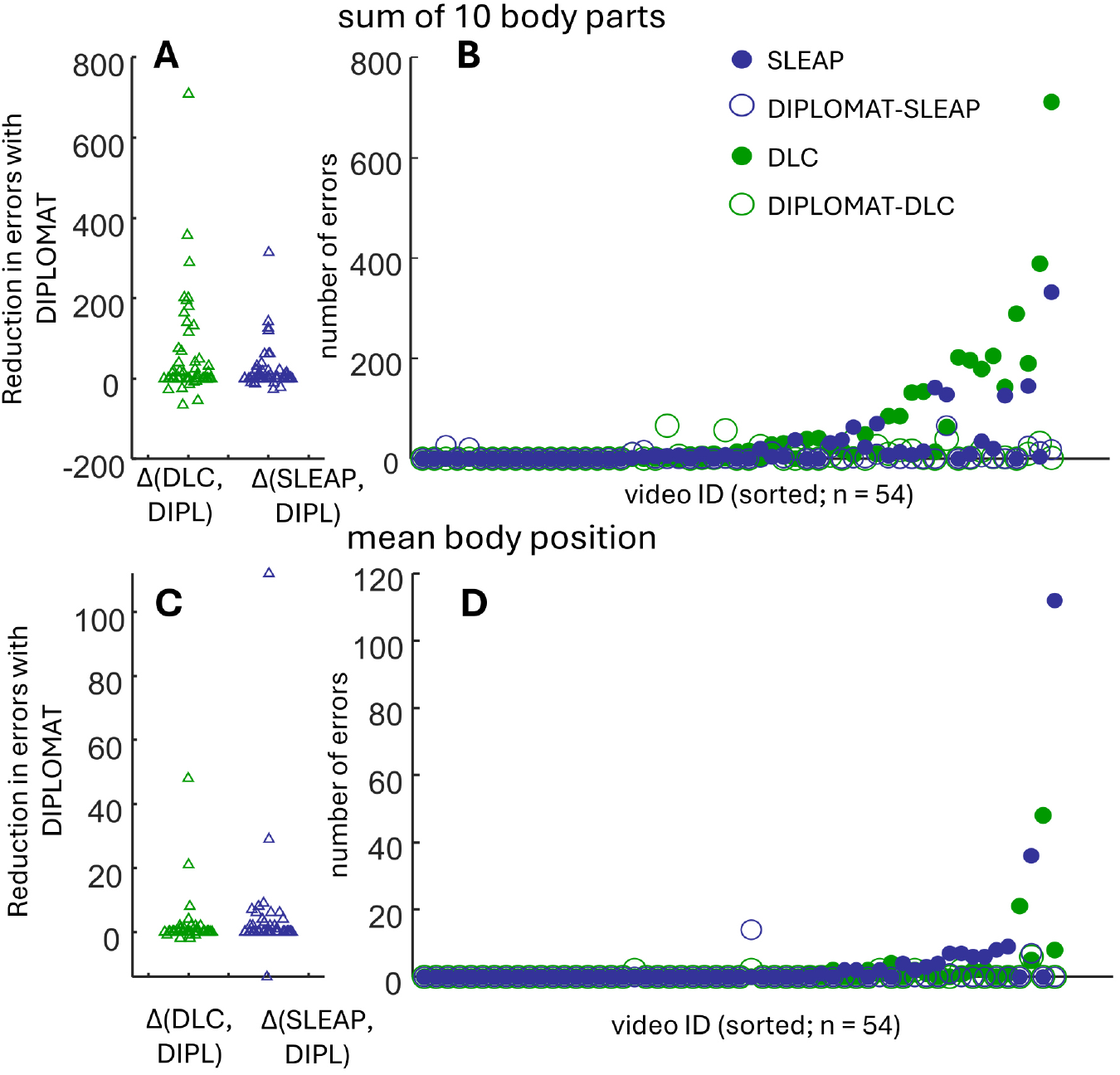
Comparison of body-swap events using the automated-only components of DIPLOMAT (Track) against automated tracking using stand-alone SLEAP or DLC. (A) Swarm plot showing the improvement, when using DIPLOMAT, in the number of cases in which any of 9 body parts across any of 3 animals switches to an incorrect individual; each point is computed for a single video by subtracting body swap errors using DIPLOMAT-DLC from DLC alone (green), and DIPLOMAT-SLEAP from SLEAP alone (blue). (B) Data across all 54 MABe triplet videos, sorted according to maximum number of swaps in either DLC or SLEAP (solid green and blue circles respectively). Although use of DIPLOMAT (open circles) occasionally resulted in more body swaps (appearing above the slope), in the majority of cases DIPLOMAT reduced or eliminated body swaps (open circles below the slope of filled circles) (C&D) Same as A&B but measuring whole-body swaps (calculated as mean body position).

#### Potential for errors

There are several ways in which Track may remain insufficient for body part location inferences. Detailed analysis of the few examples where using DIPLOMAT yielded more errors than stand-alone DLC or SLEAP (illustrated by the markers in Figure 2A&C that fall below 0) revealed that these often were videos in which two or three mice remained in physical contact throughout the entire 60 s video. When this happens, DIPLOMAT is unable to identify a strong “fix frame” (a video frame in which bodies are definitively separate). The independent-trace Viterbi is therefore initiated from frames without a meaningful prior, and identifications are more easily swapped between bodies. In theory, another way DIPLOMAT differs from status-quo methods is if a body leaves and then quickly returns to a location. In this case, it seems feasible that independent-trace Viterbi will maintain the body part locations between the movement frames, treating it as a more probable option than following the fast moving animal across frames. We have not directly observed this to happen when Track is run with default parameters. Another error case is common to all tools: when animals overlap with one-another (consequently pushing entropy of the probability distributions low) there is an increasing chance that probability fields will be split in the wrong direction, causing an identity swap. Although these challenges can be addressed with further algorithmic changes, without use of multiple cameras, it is difficult to envision any tool that perfectly tracks multiple animals without cases of identity confusion. We therefore develop a follow-up tool `Interact` that provides a memory efficient and intuitive user interface to review, edit, and re-track.

### Interact: editing tracks using the DIPLOMAT interface

We developed DIPLOMAT’s `Interact` to allow users to quickly correct errors remaining after automated analysis in the `Track` stage. The `Interact` workflow can be summarized as following three steps, often repeated across multiple cycles when reviewing a video: (1) review tracked positions, (2) manually edit body part positions, and (3) re-run automated trace routines. Figure 3 provides a labeled screenshot of the `Interact` interface.

**Fig 3.**
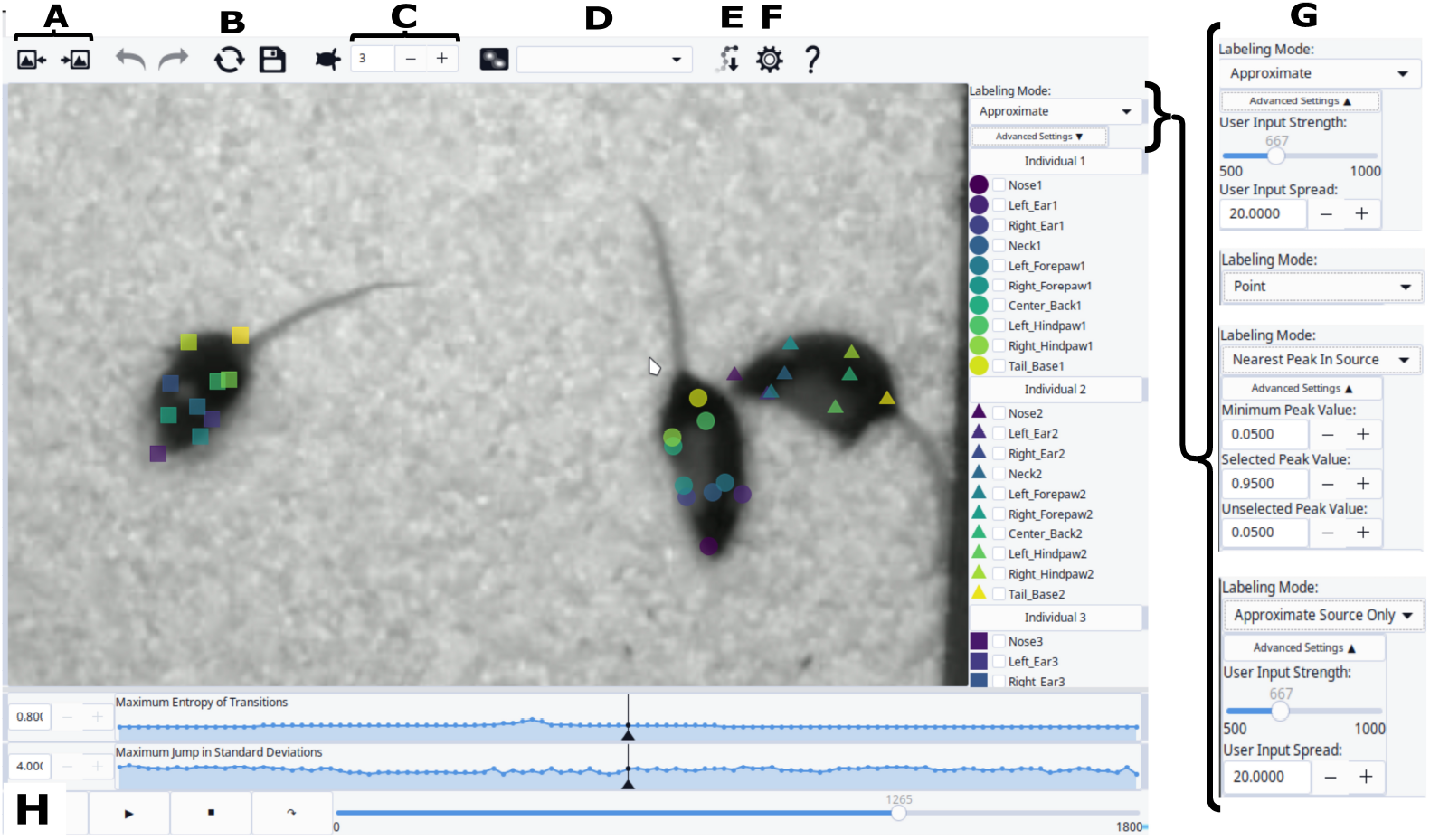
Interact can be used to edit body parts after automated tracking. Image includes letter labels associated with interface tools, as follows: (A) “Frame skip” advances to the next (or prior) section of video where tracking errors are more likely, as measured by entropy and frame-to-frame peak jumps (see H). (B) “Re-run” quickly re-applies automated tracking algorithms, incorporating the edits. (C) “Hover-track speed”: adjusts video playback speed when holding down the CTRL key to continuously label (i.e., using “hovering-track” functionality). (D) The user can optionally overlay the probability fields produces by the inference engine, perhaps to inspect the cause of labeling errors. The drop-down box allows the user to select which body part (e.g. all noses) will be shown. (E) “Export” creates a CSV file with all inferred body part positions throughout the video. (F) “Settings” allows user to adjust marker sizes and colors. (G) “Labeling mode” switches between label modes, which are optimized for different conditions (e.g., editing single versus multiple body parts, adjusting for poor probability maps, correcting identity swaps versus occlusion, etc.). Each labeling mode has adjustable parameters. (H) Plots showing frame-by-frame measures of tracking confidence, based on transition matrix entropy (related to animal proximity) and distance that peaks moved. User-defined thresholds help to flexibly identify concerning sections of the video.)

To support rapid identification of potential errors, DIPLOMAT computes two per-frame measures that may highlight frames with a high risk of error: (i) maximum transition entropy and (ii) maximum jump. Maximum entropy is a measure of the extent to which a body part instances in one frame may have more than one viable candidate for pairing in the next frame; a low value indicates that individuals are well separated, while a high value indicates that at least two animals are in close proximity. The jump distance is calculated by determining the Euclidean distance of each body part instance in the frame to its location in the next frame, then dividing that by the standard deviation of movement for that body part; the largest of these values is the *Maximum Jump in Standard Deviations*.

When using the `Interact` interface, the tracked video can be played with labels overlaid; the user can watch the entire labeled video, or may choose to skip to the next/previous point in the video with above-threshold entropy/jump values. After identifying a tracking error, the user can correct the error with either precise or broad actions. One option is to select a body part, then click to place it on the screen, optionally placing the part continuously across many frames while pressing the CTRL key. Another option is to place/correct many body parts concurrently – this is performed by selecting a set of parts (or an entire body), then clicking onto the screen. In both cases, DIPLOMAT provides a labeling mode (upper right corner of window, Figure 3), that often substantially improves speed of placement. In the “Nearest Peak in Source” labeling mode, each body part label finds the matching probability peak nearest to the user mouse placement, and assigns the part to that location. This can be particularly useful in cases where the initial automated tracking has fully swapped two bodies (i.e., the user is pulling points away from one set of local minima to another). Other labeling modes, such as “exact” and “approximate”, are better suited for single-body part labeling, such as following a misplaced body part.

After one or more frames has been edited, the user can re-run the automated tracing algorithm. Prior to re-running the Viterbi maximum trace probability algorithm, probability maps are updated to treat edited locations as 100% confident, so that the trace is forced to use that placement. As a result, it is often only necessary to edit one or a few frames to fix a body swap.

DIPLOMAT Track and Interact have been applied to multiple datasets in mice (e.g., MABe), rats (triad and dyad), and degus (triad and dyad, including with head implants, and in full darkness using an IR camera with longer exposure times). Virtually all multi-animal videos tested show cases of either body part occlusions or swaps of at least one body part; Figure 4 provides a few example comparisons between frames tracked using SLEAP versus DIPLOMAT. The key difference between manual editing using DIPLOMAT Interact compared with SLEAP is that, since DIPLOMAT preserves probability fields in its “.dipui” files used for Interact, these can help guide and facilitate editing. Current estimates of manual edit time differences between SLEAP and DIPLOMAT are roughly 75% of video length time (±30% STD). Table 3 summarizes manual editing tools available for DIPLOMAT compared with SLEAP.

**Table 3.**
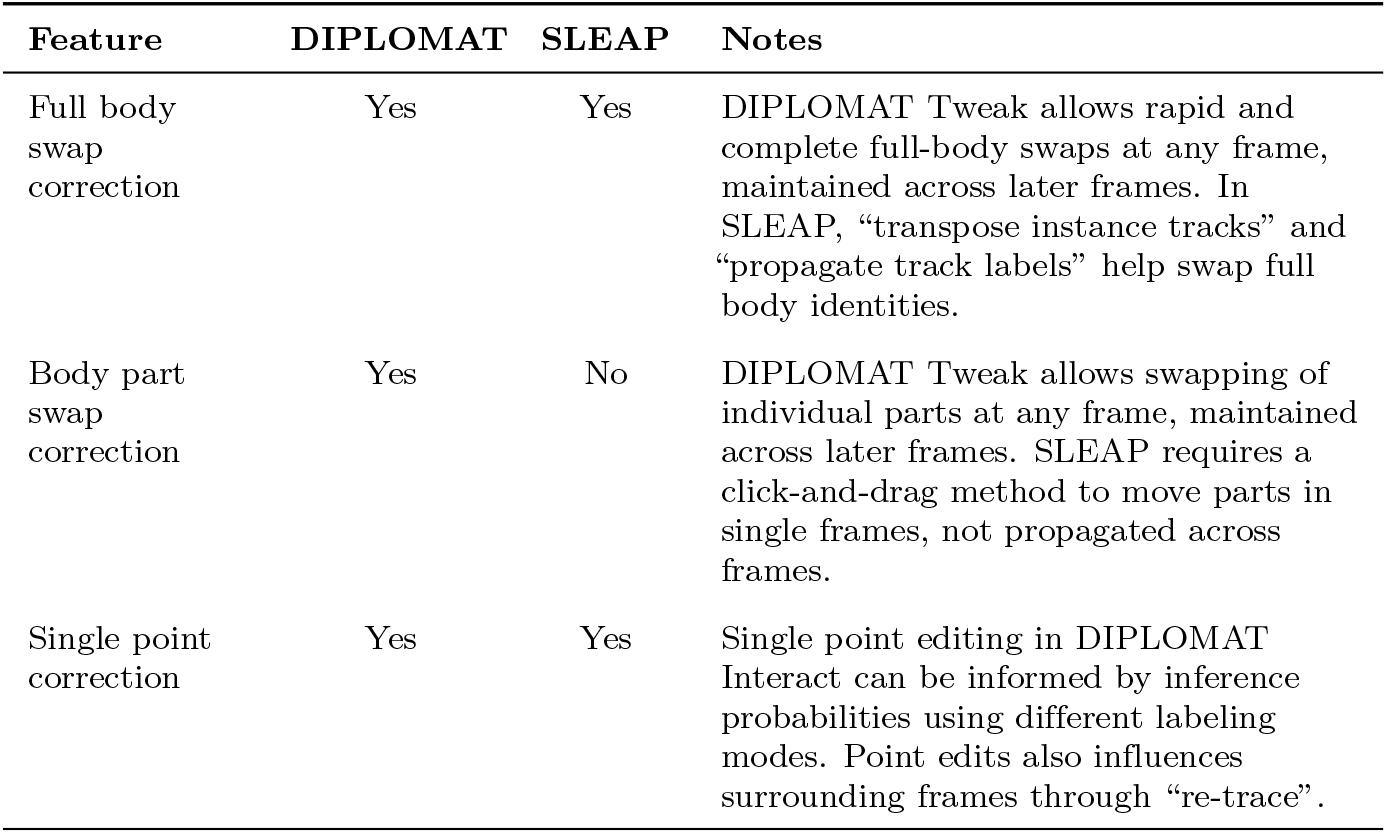
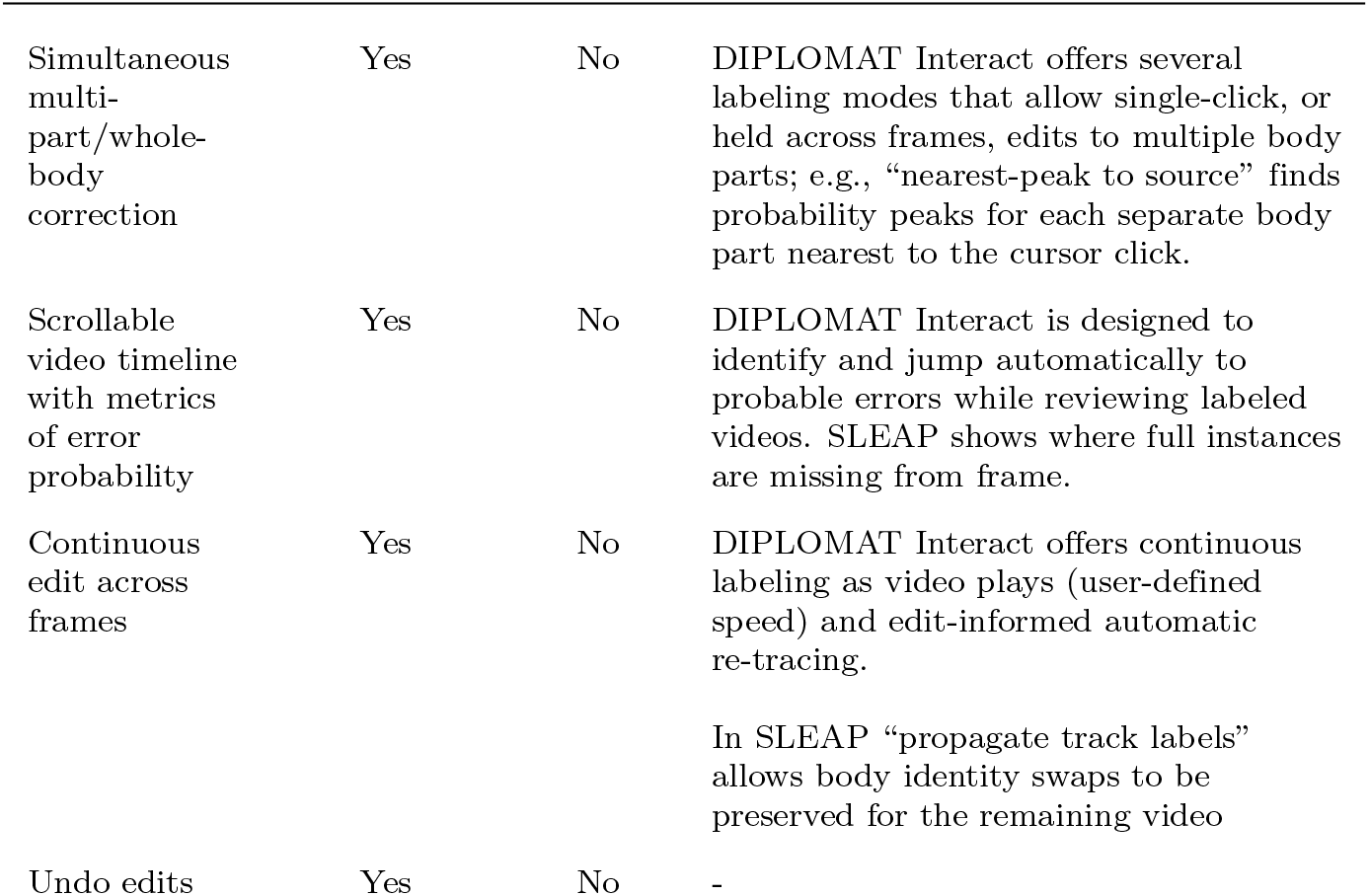
Comparison of error-correction features between DIPLOMAT and SLEAP.

| Feature | DIPLOMAT | SLEAP | Notes |
| --- | --- | --- | --- |
| Full body swap correction | Yes | Yes | DIPLOMAT Tweak allows rapid and complete full-body swaps at any frame, maintained across later frames. In SLEAP, “transpose instance tracks” and “propagate track labels” help swap full body identities. |
| Body part swap correction | Yes | No | DIPLOMAT Tweak allows swapping of individual parts at any frame, maintained across later frames. SLEAP requires a click-and-drag method to move parts in single frames, not propagated across frames. |
| Single point correction | Yes | Yes | Single point editing in DIPLOMAT Interact can be informed by inference probabilities using different labeling modes. Point edits also influences surrounding frames through “re-trace”. |
| Simultaneous multi-part/whole-body correction | Yes | No | DIPLOMAT Interact offers several labeling modes that allow single-click, or held across frames, edits to multiple body parts; e.g., “nearest-peak to source” finds probability peaks for each separate body part nearest to the cursor click. |
| Scrollable video timeline with metrics of error probability | Yes | No | DIPLOMAT Interact is designed to identify and jump automatically to probable errors while reviewing labeled videos. SLEAP shows where full instances are missing from frame. |
| Continuous edit across frames | Yes | No | DIPLOMAT Interact offers continuous labeling as video plays (user-defined speed) and edit-informed automatic re-tracing.<br>In SLEAP “propagate track labels” allows body identity swaps to be preserved for the remaining video |
| Undo edits | Yes | No | - |

**Fig 4.**
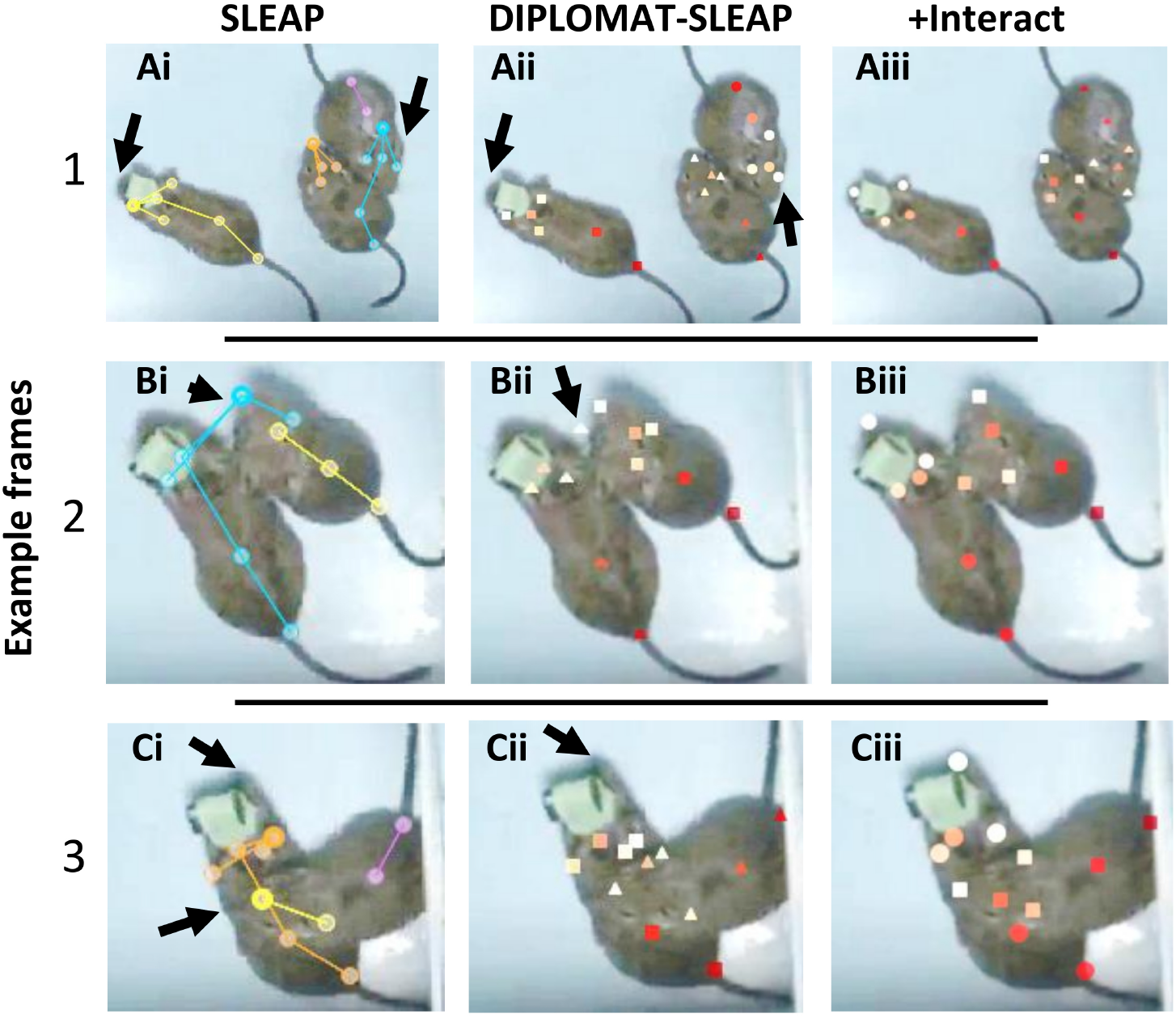
Example visual comparison between SLEAP and DIPLOMAT. Frames are taken from a single, arbitrarily-selected 5 min video with a top-down view of 3 interacting degus. All panels show example frames cropped to highlight specific errors (sometimes omitting the third degu). One degu can be easily identified by a data-logger skull implant (taped in green). Left panels (Ai, Bi, Ci) show unedited position estimates from SLEAP v1.4.1. Lines connecting markers indicate an “instance”, or inferred body skeleton. Middle panels (Aii, Bii, Cii) show position estmates from DIPLOMAT Track (v0.4.1) using the same SLEAP model. Colors are different body parts, shapes denote different identities. Right panels (Aiii, Biii, Ciii) show post-Interact edits. Black arrows point to specific errors. (Ai) A “chimera” is shown, in which body part identities (those of a single instance; blue trace) span the rear of one degu and the head, with reverse-pointing nose, of another. Four individuals are estimated, which can be merged and re-oriented in SLEAP. (Aii) DIPLOMAT-Track resolves the chimera; however, it still fails to correctly place two of the three noses. By using Interact, these can either be placed precisely (point labeling mode), snapped to the probability peak nearest a cursor-click (nearest peak labeling mode), or estimated using another labeling method (e.g., approximate labeling mode). Middle (B) and bottom (C) panels show two additional example frames with similar errors and edits.

**Fig 5.**
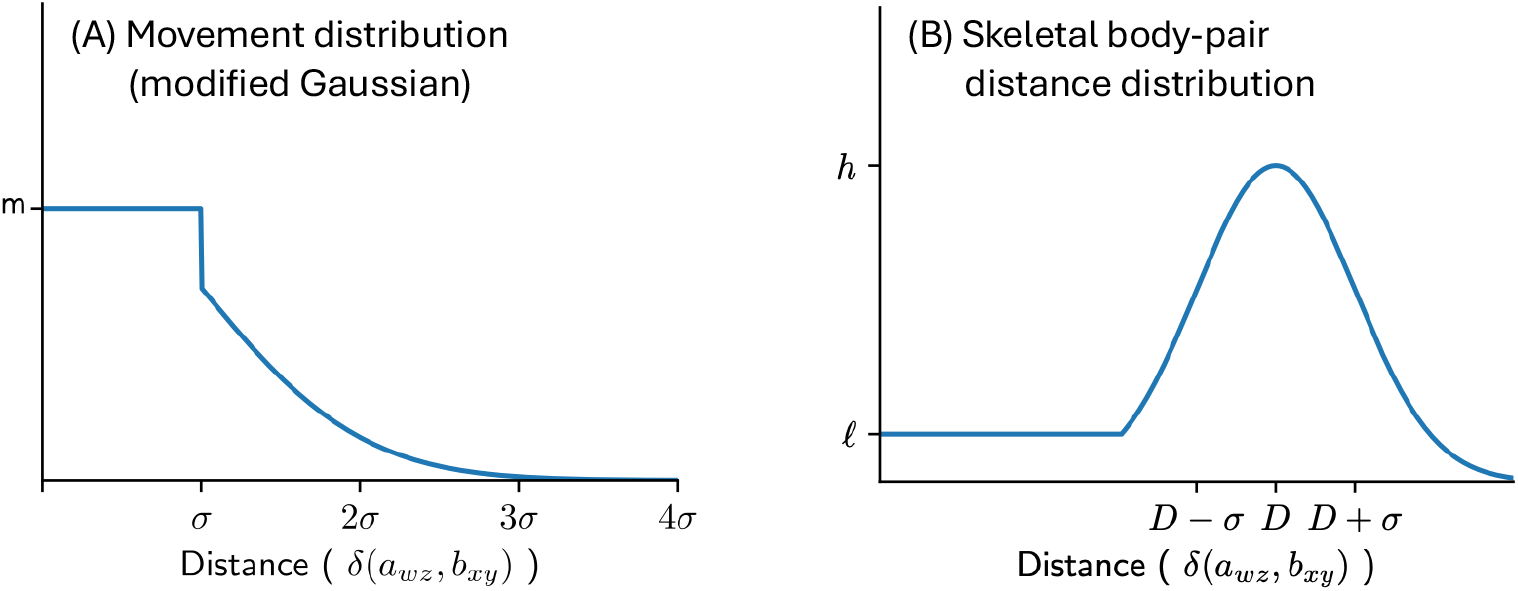
Probability density functions for (A) body part movement and (B) skeletal influence between body parts, as employed during calculation of a maximum-probability Viterbi trace. (A) The movement PDF is essentially a Gaussian, so that large movements are improbable; because small movements should not be punished relative to total lack of motion, all distances within one standard deviation of movement distance are shifted to be equally probable. (B) The skeletal PDF for a pair of body parts is designed to place high probability that one body part is the appropriate (skeletally-constrained) distance from the other; because bodies are flexible, the PDF maintains some non-zero probability even for distances much smaller than is typical.

## Discussion

The DIPLOMAT software package offers a new set of algorithms and interface tools for tracking multiple, freely moving animals in single-camera, controlled environments. We showed that these methods reduce body part and full-body swaps compared with other state-of-the-art packages, DLC and SLEAP. DIPLOMAT also produced other improvements, particularly in identity recall and, when compared against SLEAP, both precision and MOTP (i.e., tracking distance from ground truth). We further described a novel, human interface tool, `Interact`, that improves manual editing of automatically-computed traces. Interact includes the ability to dynamically re-trace (re-run the independent-trace Viterbi algorithm) across frames, thereby incorporating user-made edits.

Beyond reducing body swaps, DIPLOMAT can improve multi-animal tracking by ensuring that all body parts are accounted for. When using stand-alone convolutional neural network tools, cases of occlusion or unique body angles often lead to lost or missing body parts. The DIPLOMAT algorithms ensure object permanence, with the combination of Viterbi and body part distance (skeletal) probabilities smoothly filling gaps when the body part is not clearly visible. If desired, users can still omit these tracking points by removing “occluded state” points. While there can be risks in inferring keypoint locations that can’t be “seen” in a given video frame, the logic is consistent with unconscious inferences (to use the terminology of Helmholtz) used by all known evolved visual systems; i.e., visual knowledge is built from wedding sensory data to an internal model. Risks are also mitigated by preserving frame-by-frame probability estimates from Track. This particular “internal model” approach is different and in some ways less powerful than others that have been used; e.g., inferences based on trained mappings to 3D pose data [18]. However, they are also compatible, and could potentially be used in parallel. Furthermore, trained models depend on the training set, and may have limited external validity across animal strains and species, while algorithms like Viterbi and skeletal information are based on parameters that can be adjusted according to context.

Although our current implementation of DIPLOMAT builds on either SLEAP or DLC, the package is designed to be adaptable to any neural network based tracking tool. Any combination of algorithms that are implemented here – the independent-trace Viterbi, skeletal probability distributions, or manual edit tools – could be added as a layer on top of other tracking methods, or wrapped within higher-level, open-source and proprietary interfaces (e.g., popular packages such as Ethovision or ANY-Maze, [19–21]). It is possible, therefore, that individual tools packaged within DIPLOMAT will have value or reach beyond the software as a whole (that the whole is, for many applications, less than the sum of its parts). As an example: the Tweak interface is designed to be highly versatile, and can be easily repurposed for other tracking methods and integrated with other data streams. Tweak offers a simplified method to evaluate and edit tracks from exported CSV files, including swapping body parts or whole bodies between individuals. It is already designed to recognize and convert CSV files generated from DLC or SLEAP, but CSV conversion from other formats would be simple to implement.

Like any open-source software, the current version of DIPLOMAT can be seen as a functional snapshot, and with time and interest can be expected to evolve according to user needs. We have begun to consider several examples: First, the methods currently have no notion of size variability, which could be introduced both by modified neural network approaches as well as dynamic (potentially Markov Model informed) skeletal distances. Second, the Gaussian, spatial transition probabilities are currently static, but minor adjustments to the code could be made to fit an animal’s instantaneous velocity. Third, the code could be adjusted to work with multi-camera methods for improved, 3D posture inferences. While many research questions can be adequately addressed by 2D models, or 3D inferences using single cameras [22], there has been increasing interest in creating detailed, 3D pose estimates from multi-camera methods, including in mice [23], primates [24] and human subjects [25]. An intriguing challenge in this domain will be adapting Interact for fast, 3D-view human editing.

Together, then, DIPLOMAT both offers an important step toward more accurate and efficient, multi-animal tracking in laboratory conditions, while also providing a set of tools and map for their usage that can be repurposed across many tracking applications. Moreover, DIPLOMAT demonstrates the utility of using classical Markov models to mitigate local-optimality issues that can be pervasive when applying neural network–based approaches to time series data.

## Acknowledgments

We are grateful to Mckenna Lowrie and Angela Le for their efforts in testing DIPLOMAT on a variety of real-world datasets. We also gratefully acknowledge the computational resources and expert administration provided by the University of Montana’s Griz Shared Computing Cluster (GSCC), and the high performance computing (HPC) resources supported by the University of Arizona TRIF, UITS, and Research, Innovation, and Impact (RII) and maintained by the UofA Research Technologies department.

## Extended Methods - A Supplement

### Pose inference engines

DIPLOMAT implements pose inference for models trained using either of two popular tools, SLEAP [4] and DeepLabCut (DLC; [5]). DIPLOMAT is run on the command line by providing a trained model and a video. Because DIPLOMAT contains an independent implementation of the SLEAP/DLC inference architectures, inference with DIPLOMAT can be performed in a computer/environment that is distinct from the SLEAP/DLC training environment. DIPLOMAT’s inference engine uses the Open Neural Network Exchange (ONNX) format and is executed via ONNX Runtime.

### Initial processing

#### Sparse positional probabilities

DLC and SLEAP neural network models compute values *N* (*i, b, x, y*) corresponding to the observation of each body part type *b* at each pixel (*x, y*) of each frame *i*. The results can be interpreted as a pixel-by-pixel pseudo-probability 0.0 *≤P* (*i, b, x, y*) 1.0. *≤*

SLEAP and DLC identify peaks of *P* (*i, b, x, y*) values and assign those locations to individuals. DIPLOMAT instead uses the complete matrices to find a good trace across frames. To reduce downstream computational burden, DIPLOMAT immediately prunes low-probability pixels from consideration, retaining only the sparse set of pixels with *P* (*i, b, x, y*) *≥*1*e −*6. In each frame, sparse pixels are assigned to *n* clusters using, by deault, the complete linkage clustering method [26].

#### Estimating body sizes

DIPLOMAT tracking is based on the assumptions that (i) large changes in position between frames are rare and (ii) all parts of a body tend to remain close to each other, at relatively consistent pairwise distances. DIPLOMAT enforces these assumptions without prior knowledge of body sizes or pixel resolution by automatically estimating the size of a body and its skeletal connections. It computes the typical distance between each body-part pair as follows:

- Consider one body part estimate, *a* is at pixel (*w, z*). This can be represented as *a*_*wz*_. A second body part estimate, *b* placed at pixel (*x, y*), represented as *b*_*xy*_. The distance between these points is the Euclidean distance: 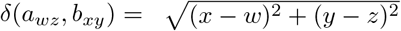.
- Each body part type *b* is represented by *n* clusters representing the *n* individuals in the video. For each body part cluster, the location is assigned to be the center of probability mass (the weighted centroid) of the cluster. Note: this positional estimate is only used for calculating inter-body-part distances, not during tracing.
- For a pair of body part types (*a, b*), we do not know in advance which cluster of *a* is correctly paired with which cluster of *b*. This makes it challenging to estimate a typical distance between *a* and *b*. DIPLOMAT performs this task by considering each instance of *a* in frame *i*, selecting the closest instance of *b* in that frame, and appending it to a vector of distances for (*a, b*); this vector eventually stores (*a, b*) distances across all frames. For each body part pair (*a, b*), DIPLOMAT then establishes estimates of the mean, *D*(*a, b*), and standard deviation, *σ*(*a, b*), for use during skeletal tracing. The median (*a, b*) distance is used as the estimated mean, and the median absolute deviation arachchige2026confidence is used as the estimated standard deviation. Empirically, the closest point is usually the correct pair, and any errors tend to slightly underestimate the skeletal distance, which is preferable for tracing.
- During tracing, DIPLOMAT uses the expected pairwise distance *D*(*a, b*) when assigning a score for placement of body parts *a* and *b* in a frame: the score is highest when the distance *δ*(*a*_*wz*_, *b*_*xy*_) between the placements is equal to *D*(*a, b*), decays to zero as the distance grows much larger than *D*(*a, b*) and decays to some constant value *>*0 for distances smaller than *D*(*a, b*). Specifically:
  – Let *h* be the maximum value assigned when *δ*(*a*_*wz*_, *b*_*xy*_) = *D*(*a, b*)
  – Let *ℓ* be the minimum value assigned when *δ*(*a*_*wz*_, *b*_*xy*_) = 0
  – Let 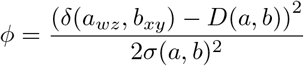

Then the skeletal value assigned to placement of parts *a*_*wz*_ and *b*_*xy*_ is:

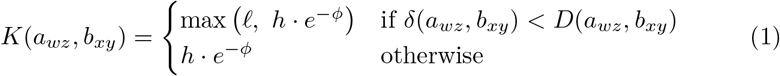

This produces a distribution of scores resembling that shown in Figure **??**B, in which a pair of body parts is most likely to have a score *h* for a distance that matches the pre-computed mode distance, decays to some constant score *ℓ* for distances as small as zero (corresponding to positioning body parts closer than is common, which can for example happen when an individual curls its body), and decays to zero for distances greater than *D*(*a, b*) (corresponding to stretching the body parts farther than they are typically observed).

Note: automatically selected skeletal lengths can be overridden by configuration settings described in the documentation.

#### Finding reliable frames and segmenting video for parallel processing

In order to improve speed, DIPLOMAT splits the video into a series of segments such that (i) a trace can be computed for each segment independently (in parallel), (ii) each segment is at least as long as a minimum segment length threshold *σ*, and (iii) the split point between two segments occurs at a frame in which individuals are well separated, so that they can be stitched back together. To achieve this, DIPLOMAT computes the reliability *R*(*f*) of every frame *f*; *R*(*f*) is a measure of the extent to which individuals are separated in the frame and subtracts the average error across skeletal connection distances. To compute separability of individuals in a frame, DIPLOMAT seeks to assign each body part to an individual in a way that maximizes the coherence of the body parts of the resulting *n* individuals. One body part group (e.g. all noses) is selected and each instance (cluster) is assigned an arbitrary identity in the range 1 to *n*. The body part group chosen as seed is the one with the largest number of skeletal connections to other body parts. When multiple parts share the same skeletal degree, the part chosen is the one that has largest minimum distance between pairs of clusters. After this, DIPLOMAT repeatedly selects the next body part *b* that minimizes the average difference between the distance of *b* and an already included part *a* compared with the expected distance *D*(*a, b*). Instances of *b* are assigned to existing individuals using the Hungarian algorithm [27] with skeletal body distance scores *K*(*a, b*) (equation 1). Once individuals have been constructed, the error across skeletal connection distances is the sum of *K*(*a, b*) values across all skeletal connections, divided by *n*.

After computing *R*(*f*) for all frames, DIPLOMAT sorts frames in descending order of reliability, adds the first frame on the sorted list to a fixed_frames list, then scans down the reliability-sorted list until reaching the next frame that is not within *σ* of an existing fixed_frame, and adds the resulting frame to the fixed_frame list. This list traversal continues until a minimum reliability threshold is reached. At this point, the list of fixed_frames acts as the list of break points between segments. In each segment, the frame *g* with highest reliability *R*(*g*) is the “anchor frame”. The purpose of selecting an anchor frame with high separability between individuals is that each individual in the frame is expected to be correctly resolved – all body parts will be correctly assigned to the same individual. This frame can serve as the starting point for a (Viterbi) maximum probability trace described below.

Within each segment, with *n* individuals, the anchor frame will contain *n* bodies (a set of body parts) assigned an arbitrary identifier between 1 and *n*. There is no reason to expect body assignment in one segment to agree with assignment in another segment, but these will be synchronized across segments in a later stage. The anchor frames are, by design, good frames for later merging of traces from consecutive segments.

### Computing a maximum probability trace

#### A hidden Markov model for multi-animal tracking

A Markov model is a statistical model that consists of a set of states and two key types of probabilities: transition (probability of moving from one state at time *i* to another state at time *i* + 1) and emission (probability that state *j* will emit some observable product at time *i*). *Hidden* Markov models (HMMs) are used to label observed sequential data that are presumed to have been emitted by the Markov model: the true path through the set of states is treated as unknown (hidden), and the task is to infer the series of states that is responsible for the observed data by computing a path through the states that has maximum probability under the transition and emission probabilities.

DIPLOMAT treats each (*x, y*) coordinate at time *i* as a state *S*(*i, x, y*). The probability that a state “emits” a specific body part *b* is taken from pseudo-probabilities produced by the pose estimation neural network: *P* (*i, b, x, y*) is the probability that body part *b* is observed at coordinate (*x, y*) and time *i*. State *S*(*i, x, y*) has non-zero transition probabilities to all states associated with time *i* + 1: (*S*(*i* + 1, *x*^′^*y*^′^) for all *x*^′^, *y*^′^), and no others; these transition probabilities follow a distance-dependent decaying function. DIPLOMAT can produce a maximum-probability trace for an individual body part through this model by following the Viterbi algorithm, in which all possible traces are considered and stored through a standard recurrence (given below), and a maximum probability path is recovered from the stored Viterbi matrix.

The between-frame movement transition probabilities are defined as a modified Gaussian distribution. The basic form of the distribution has a mean of zero and a standard deviation that is automatically determined as follows: for each instance of body part *a* in frame *i*, identify the nearest instance of *a* in frame *i* + 1 and consider that to be the matched entry; consider this to be the distance moved for that instance from frame *i*. The mean and standard deviation are computed after accumulating values over all instances of *a* and all frames in the video. This distribution is designed to allow movement between frames such that small movements are more likely than large movements, and very large movements are effectively impossible. A simple Gaussian will inappropriately penalize small movement relative to non-movement, so DIPLOMAT establishes a small radius of movement that should be treated as equally probable, computes the sum of the Gaussian probabilities of all pixel steps within that radius, then evenly distributes the sum across those pixel steps. The resulting transition probability function *T* (*x*^′^, *y*^′^, *x, y*) is a flat-topped Gaussian-like distribution such as the one in Figure **??**A.

We note that this is an atypical emission probability model in which there exist a large number of states specific to each time point, and each state’s emission probability is connected to the emission probability of each other state at the same time point. Moreover, the emission probabilities are not baked into a predefined model, but are instead the result of a complex data-dependent function: a neural network. One notable disconnect between the model and reality is that a body part can only truly appear in one position, but the model allows for the possibility that it may appear (with some probability) in many (or zero) positions. Even so, the resulting model, with defined transition probabilities and position-specific emission probabilities, is a legal HMM, and can serve as the basis of a maximum probability trace.

#### Maximum probability trace for a single individual

For a single individual *k* within a single segment starting at frame *s* and ending at frame *e*, a maximum probability trace is computed by establishing each body part *b* at the location (*x, y*) in frame *s* with the highest probability within the cluster for *b* that is assigned to individual *k*. From this start point, a probability *V*_(*i*,*b*,*k*)_(*x, y*) is computed for path for body part *b* for individual *k* running through position (*x, y*) in frame *i*, based on the recurrence:

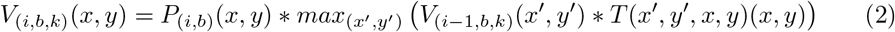

where *P*_(*i*,*b*)_(*x, y*) is the pseudo-probability (from DLC/SLEAP) of body part *b* at pos (*x, y*) in frame *i*, and *T* (*x*^′^, *y*^′^, *x, y*) is the probability of a body part moving from (*x*^′^, *y*^′^) to (*x, y*) between two consecutive frames. This recurrence is completed up to frame *e*, with a the maximum probability path recovered using the Viterbi algorithm.

By default, this simple recurrence is modified by incorporating an additional skeletal modifier that seeks to keep together all body parts for a single individual through the course of a trace. When computing the trace of a single instance of body part *b* in individual *k*, the influence in frame *i* from one other body part *a* is determined by considering all possible positions (*w, z*) of *a* for *k* in frame *i −* 1, and computing the skeletal value from equation 1. Note that the set of *K*(*a*_*wz*_, *b*_*xy*_) values is not a proper probability distribution (the values do not sum to 1.0), but can be treated as such for the purpose of computing a maximum probability trace, since values could be linearly scaled to produce normalized probabilities.

The full trace recurrence computes an *α*-weighted mixture of body movement (equation 2) and geometric mean over skeletal influences among the set *A*_*b*_ of body parts connected to *b*, where the default *α* = 0.05 moderates the skeletal influence (note: the set of skeletal connections can be configured, or can default to a completely interconnected graph):

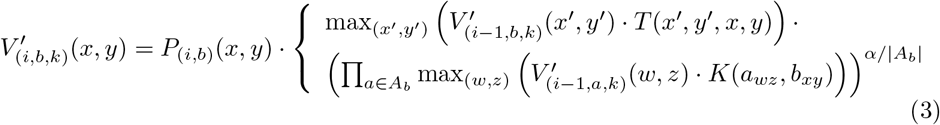

### Mutually-independent maximum probability traces

Tracking multiple individuals in a segment of the recording can be achieved by repeating the process from the previous section for each individual: for individual *k* within the segment, begin with the body part positions assigned to *k* in the anchor frame, and trace a maximum probability skeleton-adjusted path for those body parts to the end of the segment. Due to transient physical proximity of individuals and subtle flaws in the movement model, independent calculation of paths can lead to multiple paths converging, so that multiple individuals overlap on the same positions in the video. In order to force individuals to remain physically separated, DIPLOMAT modifies the Viterbi calculation to enforce mutually independent traces. For each body part type *b* and for each pixel (*x, y*) in frame *i*, DIPLOMAT identifies the instance *k* with maximum *V*_(*i*,*b*,*k*)_(*x, y*), and considers this to be the dominant individual for that pixel. This is appropriate because individual *k* will always dominate others individuals for any path leading through (*x, y*) at frame *i*. For all other individuals *h, V*_(*i*,*b*,*h*)_(*x, y*) is set to zero, to ensure that paths for those individuals do not collapse into a pixel where they are dominated by *k*.

### Occluded states

In some frames, a body part may be entirely missed by the pose prediction model. Because DIPLOMAT identifies a most probable path through only pixels with non-trivial probability, the result will be that the natural path is lost and the body part must jump to some other location. To avoid this tracing error, DIPLOMAT makes use of “occluded states”, which model the unobserved presence of a body part in a pixel. The probability held in the occluded state for body part *b* at pixel (*x, y*) in frame *i* is *O*_(*i*,*b*)_(*x, y*) = *P*_(*i*−1,*b*)_(*x, y*) *·t*_*o*_, where *t*_*o*_ is the probability of transitioning to an occluded state. Return from occluded state to visible state is possible, and adds an additional competitor *O*_(*j*−1,*b*)_(*x, y*) to the *max* function of *V*_(*j*,*b*)_(*x, y*).

### Stitching trace segments

Within each segment, the result of the above tracking steps is a collection of *n* sets of body part traces. Importantly, the identity associated with each set is arbitrary, since assignments were made on anchor frames with no communication between segments. Time-adjacent segments are connected by an anchor frame, which by design is fairly-well separable – this allows straightforward strategy for synchronizing identities across segments together. Adjacent segments are joined by considering positions of body parts in the final frame of the earlier segment and the first frame of the later segment. Typically, matching of individuals from one frame to the next is a straightforward matter of linking individuals that are closest to each other across frames, but movement can sometimes confound a greedy approach. For a more robust stitching is achieved in two steps: (1) for *n* individuals, an *n*-by-*n* distance matrix is computed, with the distance of individual *i* in the preceding frame to individual *j* in the following frame computed as the sum of body part Euclidean distances between the frames, and (2) the Hungarian assignment algorithm is applied to optimally match all individuals. Following pairwise stitching of individuals across segments, DIPLOMAT modifies all individual identities in the second segment to match the assignment from the segment containing the highest quality anchor frame.

### Interact

Because automated tracking assignments are likely to make mistakes across long video recordings, DIPLOMAT is released with a graphical user interface (GUI) that implements what we call the *Interact* stage of DIPLOMAT, and simplifies the task of manually correcting automated errors. An important feature of the GUI is its tight integration with the DIPLOMAT tracking algorithm: after modifying body-part locations on a few frames, a user may cause the Viterbi tracing algorithm to be re-run while accounting for the changes. Internally, DIPLOMAT modifies probability matrix to place 100% confidence in the manual label (i.e. when the user has placed body part *b* at position (*x, y*) of frame *i*, then *P* (*i, b, x, y*) == 1 and *P* (*i, b, w, z*) == 0 for all (*w, z*) ≠ (*x, y*)), then re-runs Viterbi. Because *P* (*i, b, x, y*) == 1, the trace must go through (*x, y*) and un-modfied errors in neighboring frames are likely to follow the corrected assignments.

Following completion of the automated trace, the *Interact* GUI helps the user to identify tracking errors by computing two measures for each frame that may indicate an increased risk of labeling errors: (1) maximum jump distance for any body part in the video (because large changes in location for an individual may indicate a body swap) and (2) a measure of the variance (entropy) of distances moved by various body parts in the video (hi entropy of movement distances suggests that one or more parts may have swapped to a new body). The user may step forward through peak values of these indicators in order to quickly find errors requiring manual intervention. Once a misplaced body part has been identified, DIPLOMAT optionally allows a mode that snaps a single body part or entire set of parts to probability peaks near the mouse-controlled position, and also allows a simple key interface to enumerate frames while controlling part position. These functions, combined with Viterbi retracing, enable faster adjustment tracking errors.

## References

1. Romero-Ferrero F, Bergomi MG, Hinz RC, Heras FJH, De Polavieja GG. Idtracker.Ai: Tracking All Individuals in Small or Large Collectives of Unmarked Animals. Nature Methods. 2019 Feb;16(2):179–82. doi:10.1038/s41592-018-0295-5.

2. Chen Z, Zhang R, Eva Zhang Y, Zhou H, Fang HS, Rock RR, et al. AlphaTracker: A Multi-Animal Tracking and Behavioral Analysis Tool; 2020. doi:10.1101/2020.12.04.405159.

3. Walter T, Couzin ID. TRex, a Fast Multi-Animal Tracking System with Markerless Identification, and 2D Estimation of Posture and Visual Fields. eLife. 2021 Feb;10:e64000. doi:10.7554/eLife.64000.

4. Pereira TD, Tabris N, Matsliah A, Turner DM, Li J, Ravindranath S, et al. SLEAP: A Deep Learning System for Multi-Animal Pose Tracking. Nature Methods. 2022 Apr;19(4):486–95. doi:10.1038/s41592-022-01426-1.

5. Lauer J, Zhou M, Ye S, Menegas W, Schneider S, Nath T, et al. Multi-Animal Pose Estimation, Identification and Tracking with DeepLabCut. Nature Methods. 2022 Apr;19(4):496–504. doi:10.1038/s41592-022-01443-0.

6. Coulombe V, Monfared S, Aghel K, Leboulleux Q, Peralta MR, Gosselin B, et al. Scaling Up Social Behavior Studies: Real-Time, Large-Scale and Prolonged Social Behavior Analysis with PrecisionTrack. Neuroscience; 2024. doi:10.1101/2024.12.26.630112.

7. Shemesh Y, Chen A. A Paradigm Shift in Translational Psychiatry through Rodent Neuroethology. Molecular Psychiatry. 2023 Mar;28(3):993–1003. doi:10.1038/s41380-022-01913-z.

8. Luxem K, Sun JJ, Bradley SP, Krishnan K, Yttri E, Zimmermann J, et al. Open-Source Tools for Behavioral Video Analysis: Setup, Methods, and Best Practices. eLife. 2023 Mar;12:e79305. doi:10.7554/eLife.79305.

9. Bordes J, Miranda L, Müller-Myhsok B, Schmidt MV. Advancing Social Behavioral Neuroscience by Integrating Ethology and Comparative Psychology Methods through Machine Learning. Neuroscience & Biobehavioral Reviews. 2023 Aug;151:105243. doi:10.1016/j.neubiorev.2023.105243.

10. Padilla-Coreano N, Batra K, Patarino M, Chen Z, Rock RR, Zhang R, et al. Cortical Ensembles Orchestrate Social Competition through Hypothalamic Outputs. Nature. 2022 Mar;603(7902):667–71. doi:10.1038/s41586-022-04507-5.

11. Kabra M, Robie AA, Rivera-Alba M, Branson S, Branson K. JAABA: Interactive Machine Learning for Automatic Annotation of Animal Behavior. Nature Methods. 2013 Jan;10(1):64–7. doi:10.1038/nmeth.2281.

12. Segalin C, Williams J, Karigo T, Hui M, Zelikowsky M, Sun JJ, et al. The Mouse Action Recognition System (MARS) Software Pipeline for Automated Analysis of Social Behaviors in Mice. eLife. 2021 Nov;10:e63720. doi:10.7554/eLife.63720.

13. Weinreb C, Pearl J, Lin S, Osman MAM, Zhang L, Annapragada S, et al. Keypoint-MoSeq: Parsing Behavior by Linking Point Tracking to Pose Dynamics; 2023. doi:10.1101/2023.03.16.532307.

14. Goodwin NL, Choong JJ, Hwang S, Pitts K, Bloom L, Islam A, et al. Simple Behavioral Analysis (SimBA) as a Platform for Explainable Machine Learning in Behavioral Neuroscience. Nature Neuroscience. 2024 Jul;27(7):1411–24. doi:10.1038/s41593-024-01649-9.

15. Forney GD. The Viterbi Algorithm. Proceedings of the IEEE. 1973;61(3):268–78. doi:10.1109/PROC.1973.9030.

16. Sun JJ, Marks M, Ulmer A, Chakraborty D, Geuther B, Hayes E, et al. MABe22: A Multi-Species Multi-Task Benchmark for Learned Representations of Behavior. arXiv. 2022. doi:10.48550/ARXIV.2207.10553.

17. Heindl C. py-motmetrics: Python implementation of multiple-object tracking metrics; 2017. Accessed: 2025-08-11. https://github.com/cheind/py-motmetrics.

18. Dunn TW, Marshall JD, Severson KS, Aldarondo DE, Hildebrand DGC, Chettih SN, et al. Geometric Deep Learning Enables 3D Kinematic Profiling across Species and Environments. Nature Methods. 2021 May;18(5):564–73. doi:10.1038/s41592-021-01106-6.

19. Noldus LPJJ, Spink AJ, Tegelenbosch RAJ. EthoVision: A Versatile Video Tracking System for Automation of Behavioral Experiments. Behavior Research Methods, Instruments, & Computers. 2001 Aug;33(3):398–414. doi:10.3758/BF03195394.

20. Spink AJ, Tegelenbosch RAJ, Buma MOS, Noldus LPJJ. The EthoVision Video Tracking System—A Tool for Behavioral Phenotyping of Transgenic Mice. Physiology & Behavior. 2001 Aug;73(5):731–44. doi:10.1016/S0031-9384(01)00530-3.

21. Lim CJM, Platt B, Janhunen SK, Riedel G. Comparison of Automated Video Tracking Systems in the Open Field Test: ANY-Maze versus EthoVision XT. Journal of Neuroscience Methods. 2023 Sep;397:109940. doi:10.1016/j.jneumeth.2023.109940.

22. Karashchuk P, Rupp KL, Dickinson ES, Walling-Bell S, Sanders E, Azim E, et al. Anipose: A Toolkit for Robust Markerless 3D Pose Estimation. Cell Reports. 2021 Sep;36(13):109730. doi:10.1016/j.celrep.2021.109730.

23. Klibaite U, Li T, Aldarondo D, Akoad JF, Ö lveczky BP, Dunn TW. Mapping the Landscape of Social Behavior. Cell. 2025 Apr;188(8):2249–66.e23. doi:10.1016/j.cell.2025.01.044.

24. Hayden BY, Park HS, Zimmermann J. Automated Pose Estimation in Primates. American Journal of Primatology. 2022 Oct;84(10):e23348. doi:10.1002/ajp.23348.

25. Wang J, Tan S, Zhen X, Xu S, Zheng F, He Z, et al. Deep 3D Human Pose Estimation: A Review. Computer Vision and Image Understanding. 2021 Sep;210:103225. doi:10.1016/j.cviu.2021.103225.

26. Sorensen T. A method of establishing groups of equal amplitude in plant sociology based on similarity of species content and its application to analyses of the vegetation on Danish commons. Biologiske skrifter. 1948;5:1–34.

27. Kuhn HW. The Hungarian method for the assignment problem. Naval research logistics quarterly. 1955;2(1-2):83–97.

